# Single cell RNA-sequencing suggests a novel lipid-associated mast cell population following weight cycling

**DOI:** 10.1101/2023.11.12.566786

**Authors:** Munira Kapadia, Alexa M. Betjemann, Matthew A. Cottam, Mona Mashayekhi, Heidi J Silver, Alyssa H. Hasty, Heather L. Caslin

## Abstract

We previously demonstrated that weight cycled mice have increased adipose mast cells compared to obese mice by single cell RNA-sequencing. Here, we aimed to confirm and elucidate these changes. Interestingly, we did not detect an increase in total mast cell numbers in weight cycled mice by Toluidine blue or flow cytometry, however, further subcluster analysis of our dataset showed that our initial mast cell cluster consisted of two unique populations. One population had very high expression of classical mast cell markers and another had elevated lipid handling and antigen presentation genes with a concomitant reduction in classical mast cell genes. This new “lipid-associated” mast cell cluster accounted for most of the mast cells in the weight cycled group. We induced a similar phenotype *in vitro* using repeated exposure to adipose tissue conditioned media to mimic weight gain and weight regain. Upon repeated exposure to adipose tissue conditioned media, bone marrow-derived mast cells had increased lipid droplets and reduced expression of cKit and FcεR1 compared to control cells. Moreover, we analyzed mast cells in a pilot study of subcutaneous adipose tissue from four obese, prediabetic women. We found two mast cell populations that appear similar to the murine populations detected by sequencing. The population with reduced cKit and FcεR1 was significantly correlated with weight variance. Together, these data suggest that weight cycling may induce a unique population of mast cells similar to lipid-associated macrophages, which have been shown to play a role in diverse diseases from obesity and atherosclerosis to Alzheimer’s disease. Future studies will focus on isolation of these cells from mice and humans to better determine their lineage, differentiation, and functional roles.

## 1. Introduction

Risk for cardiometabolic diseases is higher with excess adipose tissue, and thus, weight loss is often prescribed. Attempts at weight loss are common, and the most recent National Health and Nutrition Examination Survey (NHANES) data indicates that ∼50% of adults in the United States attempted weight loss between 2013-2016(1). Other reports suggest that up to 76% of adults in the United States have attempted weight loss at some point in their lifetime(2). Weight regain is also a common occurrence, and most individuals regain most of the lost weight within just a few years(3–8). This process of repeatedly gaining and losing weight, termed weight cycling, further increases the risk for type 2 diabetes, cardiovascular disease, and hypertension beyond that of stable long-term obesity(9–13).

During weight gain, and specifically adipose expansion, immune cell recruitment to the adipose tissue promotes adipocyte lipolysis, fibrosis, and insulin resistance(14). These changes contribute to the development of type 2 diabetes and other cardiometabolic diseases. With adipose expansion, type 1 immune subsets like T helper 1 cells, CD8^+^ T cells, and neutrophils dominate, and macrophages shift from a tissue regulatory phenotype to a pro-inflammatory phenotype that induces insulin resistance and promotes ectopic fat deposition(15,16). There is also an increase in lipid-associated macrophages that help buffer excess lipid that restrain weight gain and glucose impairments(17,18). However, less is known about how immune cells change and contribute to systemic metabolism with weight cycling. We recently used single-cell RNA-sequencing (scRNAseq) to classify differences in proportion and phenotype of gonadal adipose tissue immune cells from lean, obese, weight loss, and weight cycled mice(19). As previously published(20), weight cycled mice had worsened glucose tolerance compared with their weight stable obese counterparts. Unexpectedly, these data also showed that weight cycling substantially increased the proportion of adipose mast cells.

Mast cells are tissue-resident cells in connective and mucosal tissues around the body(21). Primarily considered type 2 immune cells, mast cells are well-studied for their protective role against extracellular parasites and their detrimental role in allergic disease and asthma(22,23). However, mast cells also have many physiological functions in the defense against bacteria and viruses(24,25) and contribute to autoimmunity and atherosclerosis(26). While mast cells have been shown to increase with weight gain, their role in adipose tissue homeostasis and expansion is not well understood, and there are contradictory results in the literature due to the use of different mast cell knockout models and different concentrations of mast cell stabilizers(27–31). Therefore, the purpose of this study was to elucidate how adipose mast cells change under the conditions of weight gain, weight loss, and weight cycling. Unexpectedly, we found a novel lipid handling phenotype in mast cells from weight cycled mice that seemingly corresponds with a human adipose mast cell population and that we can replicate in culture.

## 2. Methods

### 2.1 Animals

Male and female C57BL/6J mice were purchased from Jackson Labs (Bar Harbor, ME, #000664). For *in vitro* experiments, mice were euthanized between 8-16 weeks of age. For *in vivo* experiments, male C57BL/6J mice were placed on 9-week cycles of 60% high fat diet (Research Diets, New Brunswick, NJ, #D12492, 5.21 kcal/g food) or nutrient-matched 10% low fat diet (Research Diets #D12450B, 3.82 kcal/g) at 8 or 9 weeks of age for a total of 27 weeks as published(19,20) and as visualized in **Figure 1A**. These diets contained the same purified ingredients and included the same relative amount of protein, fiber, and other micronutrients. Food and water were provided *ad libitum*, and body weight and food intake were recorded weekly.

**Figure 1:**
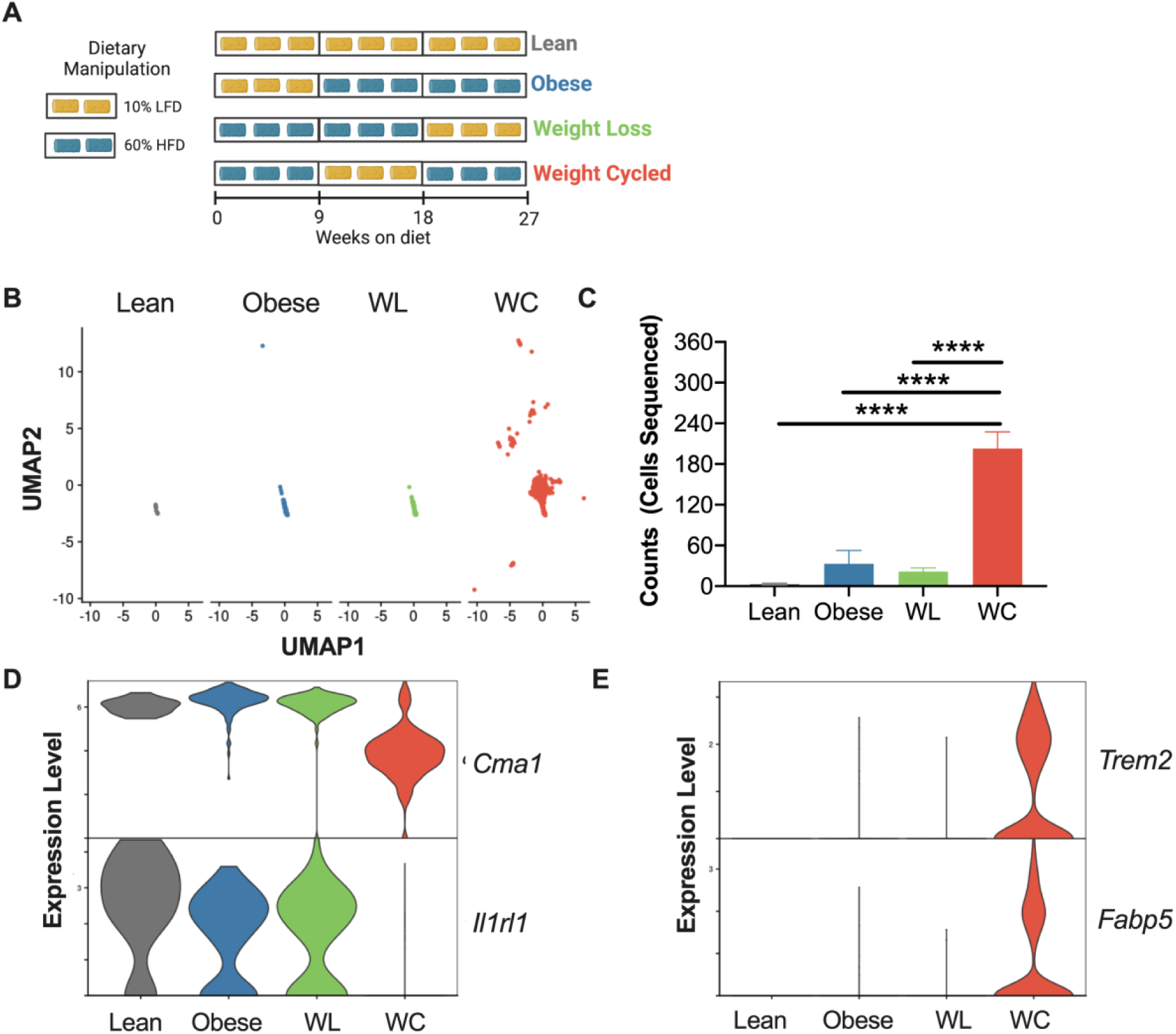
Weight cycling increases adipose mast cells. (A) Schematic of weight cycling mouse model commonly used by our group. (B) UMAP of mast cell cluster in each diet group. (C) Mast cell counts (number of cells sequenced). Violin plots of (D) *Cma1* and *Il1r1* and (E) *Trem2* and *Fabp5* expression in each diet group. All data were derived from new analyses of our previously published dataset [17].

After 26 weeks on diet, mouse body fat and muscle mass were measured by nuclear magnetic resonance whole body composition analysis using a Minispec (Bruker, Billerica, MA). The following day, mice were fasted for 5-h and briefly anesthetized with isoflurane to obtain blood via tail snip. After a 1-h recovery, basal blood glucose levels were measured using a hand-held glucometer (Contour Next EZ, Bayer, Pittsburg, PA). Mice were injected intraperitoneal with 1.5 g dextrose/kg lean mass and blood glucose was measured at 15, 30, 45, 60, 90, and 120 min.

All murine procedures were approved and performed in compliance with the Vanderbilt University or the University of Houston Institutional Animal Care and Use Committees. Both Vanderbilt University and the University of Houston are accredited by the Association for Assessment and Accreditation of Laboratory Animal Care International.

### 2.2 Single cell analysis

We used our previously published single cell sequencing dataset to further assess adipose mast cells (GEO accession number GSE182233)(19). In this previous dataset, all cluster annotations were performed using common cell type markers and confirmed via SingleR V1.6. Here, we subset the data on what was initially clustered as mast cells and performed further clustering using the FindClusters function in Seurat V4(32) with a resolution of 0.1. Differential expression analysis was performed in R version 4.1 with Seurat V4 using Wilcoxon Ranked Sum tests with the FindMarkers function to compare the new subclusters. GO Biological Processes pathway analysis was conducted on differential expression with ShinyGO 0.76.2(33) using FDR cutoff of 0.05 (last date of access 10/17/2022).

### 2.3 Histology and toluidine blue staining

Epididymal adipose tissue was fixed in 4% paraformaldehyde for 24-h, then washed with phosphate buffered saline (PBS). Samples were embedded in paraffin and then sectioned with the help of the Translational Pathology Shared Resource core at Vanderbilt University. Slides were the deparaffinized in xylene and rehydrated, then stained in 10% Toluidine blue for 5 min. After staining, slides were dehydrated, cleared in xylene, and mounted. Slides were imaged with an inverted DMi8 microscope and Flexacam color camera (Leica, Teaneck, NJ) at 20x. Multiple images were taken using the tilescan function to collect 104 sections/image. Mast cells were counted per 104 frames for two tissue sections per sample and counts for each sample were averaged.

### 2.4 RNA isolation and real time PCR

RNA was isolated from subcutaneous adipose tissue with a Qiagen RNeasy Mini Kit (Qiagen, #74104) according to manufacturer specifications. Following quantification with a Nanodrop, the RNA was converted to cDNA using iScript Reverse Transcription Supermix for RT-qPCR (Bio-rad, Hercules, CA, #1708840) or Sensifast cDNA Synthesis kit (Meridian Biosciences, Memphis, TN, #BIO-65054). Real Time PCR was completed using Taqman primer-probe sets from Thermofischer and iQ Supermix (Bio-Rad #1708860). Gene expression was normalized via ΔΔCt method to a housekeeping gene and the lean control.

### 2.5 Murine adipose immune cell isolation

Epididymal and subcutaneous adipose tissue were collected as previously described(34). Following isoflurane overdose and cervical dislocation, mice were perfused with 20 ml PBS through the left ventricle. Adipose fat pads were collected, minced, and digested in 6 ml of 2 mg/mL type IV collagenase (Worthington Biomedical, Lakewood, NJ, #LS004186) for 30 min at 37°C. The collagenase was diluted with cold PBS and digested tissue was vortexed and filtered through 100 μm filters. The pelleted stromal vascular fraction was then lysed with ACK buffer to remove red blood cells and filtered through a 35 μm filter to produce a single cell suspension.

### 2.6 Bone marrow-derived mast cell experiments

Bone marrow was isolated from femurs of male and female C57BL/6J mice between 8-16 weeks of age. Bone marrow-derived mast cells were (BMMC) were differentiated over 4 weeks in complete RPMI (with 10% FBS, 1% Penicillin/streptomycin, 1% HEPES, and 1% sodium pyruvate) with 5 ng/mL mouse IL-3 and SCF (Fujifilm/ Shenandoah Biotech # 200-01 and 200-09). Differentiation was confirmed by flow cytometry for cKit and FcεR1 and cells were used once >90% mast cell differentiation was achieved.

For adipose tissue conditioned media (ATCM), male C57BL/6J mice were purchased from Jackson labs on high fat diet (DIO; #:380050) or fed 60% high fat diet (Research Diets #D12492) for 9-12 weeks. Mice were euthanized and epididymal adipose fat pads were collected in cold PBS. In sterile conditions, adipose fat pads were cut into small pieces, washed in PBS, and then cultured in complete RPMI for 48 hours at ∼50 mg/mL. Media was then centrifuged to remove the top adipocyte fraction and the pelleted stromal vascular fraction and stored at −20°C for further experiments.

For lipid handling induction, BMMC were treated ± ATCM for 24 hours, replated in standard RPMI media for 3 days, and treated again ± ATCM for 24 hours. Mouse IL-3 and SCF (5 ng/mL) were included in all conditions throughout the culture period. Surface proteins and Bodipy staining, were analyzed by flow cytometry.

### 2.7 Human data and adipose samples

Human subcutaneous adipose tissue samples were collected from an ongoing clinical study (NCT04907214). The study was approved by the Vanderbilt Institutional Review Board and conducted according to the Declaration of Helsinki. All participants provided written informed consent. We obtained adipose samples paired with deidentified demographic information and self-reported weight history from four participants. Inclusion criteria include women with obesity (BMI ≥ 30kg/m^2^) and pre-diabetes (fasting glucose of 100-125mg/dL, or two-hour plasma glucose of 140-199 mg/dL during a 75-gram oral glucose tolerance test, or hemoglobin A1c of 5.7-6.4%). Additionally, all participants completed a weight history questionnaire asking about weight over the past 6 months, one year, two years, five years, and ten years along with any noted diet attempts at weight loss. As an indicator of weight cycling, we calculated sample weight variance (standard deviation squared) for each subject.

Participants underwent bedside liposuction for collection of adipose tissue from the periumbilical region using a Tulip^TM^ Medical closed syringe system (Tulip Medical Products, San Diego, CA). Under aseptic conditions and local lidocaine anesthesia, a small incision was made in the skin. The GEMS Johnnie Snap lock was placed inside a 60-cc syringe with the Tulip liposuction cannula attached and was inserted at an angle through the incision to below Scarpa’s fascia. Suction was applied until the syringe activated the clicker lock. The needle was oscillated at a rate of approximately 1 Hz without breaking suction with a twisting motion. The sampling continued until 1-10 grams of tissue was removed. The tissue sample was placed in 20-30 cc of cold saline and immediately processed for enrichment of the stromal vascular fraction.

Harvested adipose tissue was rinsed with PBS and weighed, then digested in gentleMACS C tubes with a Dissociator (Miltenyi Biotec, Gathersburg, MD). Collagenase D (Roche, Indianapolis, IN) was added at 2 mg/mL and the sample was shaken at 37 °C degrees at 150 rpm for 40 min. The digested tissue was rinsed, filtered through a 70-micron filter, and the floating adipocyte fraction removed. The remaining immune-rich stromal vascular fraction was pelleted and washed, then used for staining for flow cytometry.

### 2.8 Flow cytometry

Murine stromal vascular cells or BMMC were prepared for flow cytometry. FcBlock (1:200) was added for 10 min. Surface proteins were stained with fluorescent antibodies at 1:200 for 30 min at 4 °C using the antibody panels in **Supplemental Table 1**. For lipid droplet staining, Bodipy was added at 1μM for 30 min at the same time as surface antibodies. All samples were washed after antibody and Bodipy staining and stained with DAPI (1:5000) immediately prior to acquisition. Data was acquired on a MACSQuant10 (Miltenyi) or 3 laser Fortessa (BD Biosciences, San Jose, CA) and analyzed on FlowJo using single color compensation controls for all stains and fluorescence minus one (FMO) controls for gating on *ex vivo* samples.

### 2.9 Statistical analyses

Statistical analyses were performed using GraphPad Prism. Student’s *t*-tests were run for comparisons between two groups, one-way analysis of variances (ANOVA)s were used for comparisons between more than two groups, and two-way ANOVAs were conducted for >2 groups over >2 timepoints. For significant main effects, *post-hoc* pairwise comparisons using Tukey or Sidak corrections were used to determine statistical differences. Simple regression was used for correlations. All data are presented as mean ± standard error of the mean (SEM). A *p*-value (or adjusted *p* - value) of <0.05 was used to determine significance.

## 3. Results

### 3.1 scRNAseq data show increased murine adipose mast cells with weight cycling

Adipose immune cell populations change with weight gain; however, less is known about how these populations change with weight loss and weight cycling. Therefore, to examine changes in adipose immune populations with weight loss and weight cycling, we performed scRNAseq in a mouse model of weight cycling and published the initial findings broadly focused on T cells, dendritic cells, and macrophages. Male mice were placed on low-fat or high-fat diets for 27 weeks total as shown in **Figure 1A**. Strikingly, in our scRNAseq study, we found that mast cell counts were significantly elevated in the weight cycled mice compared to all other diet conditions(19), and for which we provide new analyses here (**Figure 1B & C**). Additionally, our previous report showed that classical mast cell genes such as chymase (*Cma1*) and the IL-33 receptor (*Il1r1*) were reduced in the weight cycled cluster while lipid handling genes such as *Trem2* and *Fabp5* were increased in the weight cycled cluster ((19) and **Figure 1D & E**).

### 3.2 Standard techniques do not identify increased mast cells in weight cycling

To confirm the previous scRNAseq data, we attempted to detect mast cells using complementary techniques in a new cohort of mice. As seen previously in our sequencing as well as other published studies(19,20,35,36), glucose clearance was significantly lower in the weight cycled group compared with the obese group during an intraperitoneal glucose tolerance test, despite similar body weight at the end of 27 weeks (**Figure 2A & B**). For initial mast cell analysis, fixed sections of epididymal adipose tissue were stained with toluidine blue, a cationic dye that stains nuclei pink and mast cells dark purple (example image shown in **Figure 2C**). Supporting the sequencing data, there was a significant increase in mast cells in the obese and weight loss groups; however, the weight cycled group did not show increased mast cells compared with any other group (**Figure 2D**). Flow cytometry was next used to quantify mast cells in epididymal and subcutaneous adipose tissue by the expression of canonical mast cell markers, cKit and FcεR1 (**Figure 2E & F**). While there was an increase in the percent and number of mast cells/gram epididymal fat in weight cycled animals compared to the lean mice, weight loss animals showed the highest number of mast cells (**Figure 2F**). We also analyzed the expression of ST2, the IL-33 receptor, and CD107a and CD63 (LAMP-3), two lysosomal proteins which can appear on the surface of mast cells following degranulation. We detected an increase in CD63 in the obese and weight cycled groups compared to the lean and weight loss groups (**Figure Supplement 1**), which could suggest changes to cellular functions like degranulation separate from changes (or lack thereof) in cell number.

**Figure 2:**
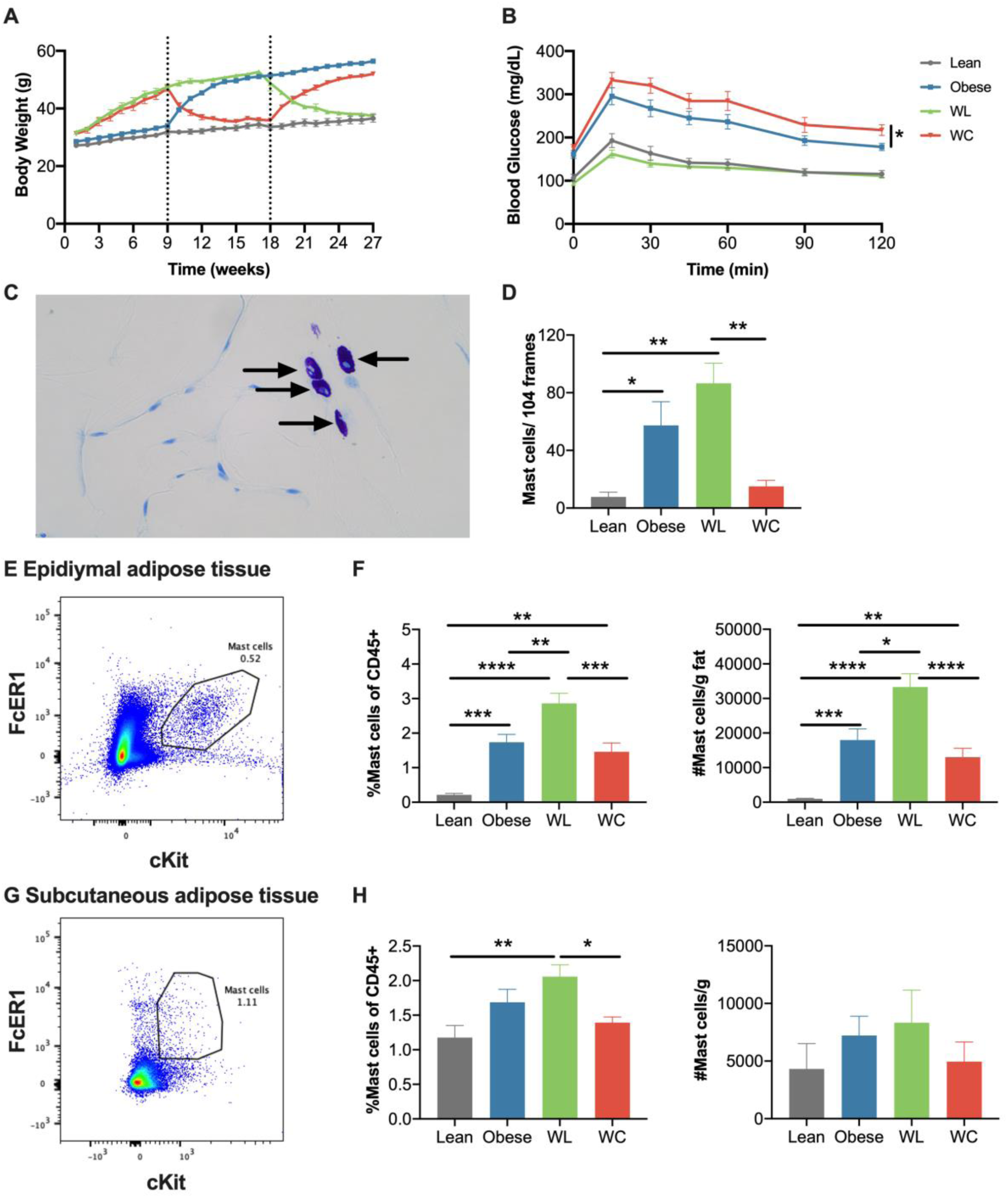
Histology and flow cytometry do not confirm an increase in adipose mast cells with weight cycling. (A) Body weight over time from a new murine cohort measured weekly with diet switch indicated by dashed lines in lean, obese, weight loss (WL), and weight cycled (WC) mice (n=12/group). (B) Blood glucose during an intraperitoneal glucose tolerance test (1.5 g dextrose/kg lean mass) at 27 weeks (n=12/group). (C) Representative image of Toluidine blue staining on adipose mast cell section. (D) Quantification of adipose mast cells per 104 frames (n=4-6/group). (E) Representative flow cytometry plot of cKit+Fcεr1+ mast cells in epididymal adipose tissue. (F) Quantification of epididymal adipose tissue mast cells by % and #/ gram fat (n=14-18/group). (G) Representative flow cytometry plot of cKit+Fcer1+ mast cells in subcutaneous adipose tissue. (H) Quantification of subcutaneous adipose tissue mast cells by % and #/g fat (n=8-12/group). Data are means ± SEM in 1-2 new cohorts of mice. *p<0.05, **p < 0.01, *** p <0.001, ****p < 0.0001.

Similar to the epididymal adipose tissue, there was a significant increase in mast cells in the subcutaneous adipose tissue of the weight loss animals compared with the lean and weight cycled groups by flow cytometry (**Figure 2G & H**). However, the number of mast cells/gram fat was more variable in this fat pad, and there were no significant differences. Real time PCR analysis was also performed on whole subcutaneous adipose tissue for *Fcer1* (the gene for FcεR1), the proteases *Mcpt4, Tpsb2,* and *Cma1 (*found in mast cell granules) and the tryptophan hydroxylase gene *Tph1 (*shown in adipose mast cells to be important for the metabolic adaptations to thermoneutrality(37)). *Fcer1* was significantly elevated in both the obese and weight cycled groups compared to the lean animals (**Figure Supplement 2**). Weight loss increased *Mcpt4, Tpsb2, Cma1,* and *Tph1*, but this was only significant compared to the lean group for *Tpsb2* and Tph1 (**Figure Supplement 2**). These results suggest that we cannot detect the increase in adipose mast cells during weight cycling by standard methods of detection.

### 3.3 scRNAseq suggests weight cycling induces a distinct cluster of lipid-associated mast cells

Toluidine blue staining and surface staining by flow cytometry are limited in that they only detect mast cell number by the presence of granules and standard surface markers, respectively. Thus, we returned to our published scRNAseq data to look more closely at transcriptional differences. First, we subset the data on the initial “mast cell” cluster which expresses high levels of mast cell specific genes (*Mcpt4*, *Cma1*, and *Tpsb2*) when compared to all other cells. Then, we recalculated the highly variable genes among the remaining cells and clustered the cells using a new uniform manifold approximation and projection (UMAP) based on the first 30 principal components. Following reclustering using the Louvain implementation in Seurat, two distinct subclusters were identified (**Figure 3A**). There was a significant increase only in cluster 2 in the weight cycled group compared with the other diet groups (**Figure 3B**). Pathway analysis of differentially expressed genes in cluster 2 compared to cluster 1 suggested a reduction in many leukocyte differentiation, hematopoiesis, and activation genes and an increase in biosynthetic and metabolic genes (**Figure Supplement 3**). Moreover, there was a significant decrease in many surface and protease genes specifically related to mast cells (**Figure 3C**). It’s important to note that mast cell genes were still increased in cluster 2 relative to other populations in our data, confirming cluster 2 is a mast cell population. We show this relative to all macrophage populations (**Figure Supplement 4**). Additionally, there was a significant increase in genes involved in lipid handling in cluster 2 as compared to cluster 1 (**Figure 3D**). Other lysosomal, scavenger, phagocytic, and antigen presentation proteins also increase in cluster 2 compared with cluster 1 (**Figure 3E, Table Supplement 2, and Figure Supplement 4**). All significant differences between clusters are reported in Supplemental Table 2. Markedly, these cells appear very similar to lipid associated macrophages, first reported in 2019(17), though they are a unique population by sequencing (**Figure Supplement 4A**), do still express some mast cell specific genes (**Supplemental Figure 4B**), and don’t express all lysosomal and scavenger genes expressed by macrophages (**Figure Supplement 4D & E**). Together, these new analyses suggest that weight cycling specifically increases a novel population of lipid handling mast cells.

**Figure 3:**
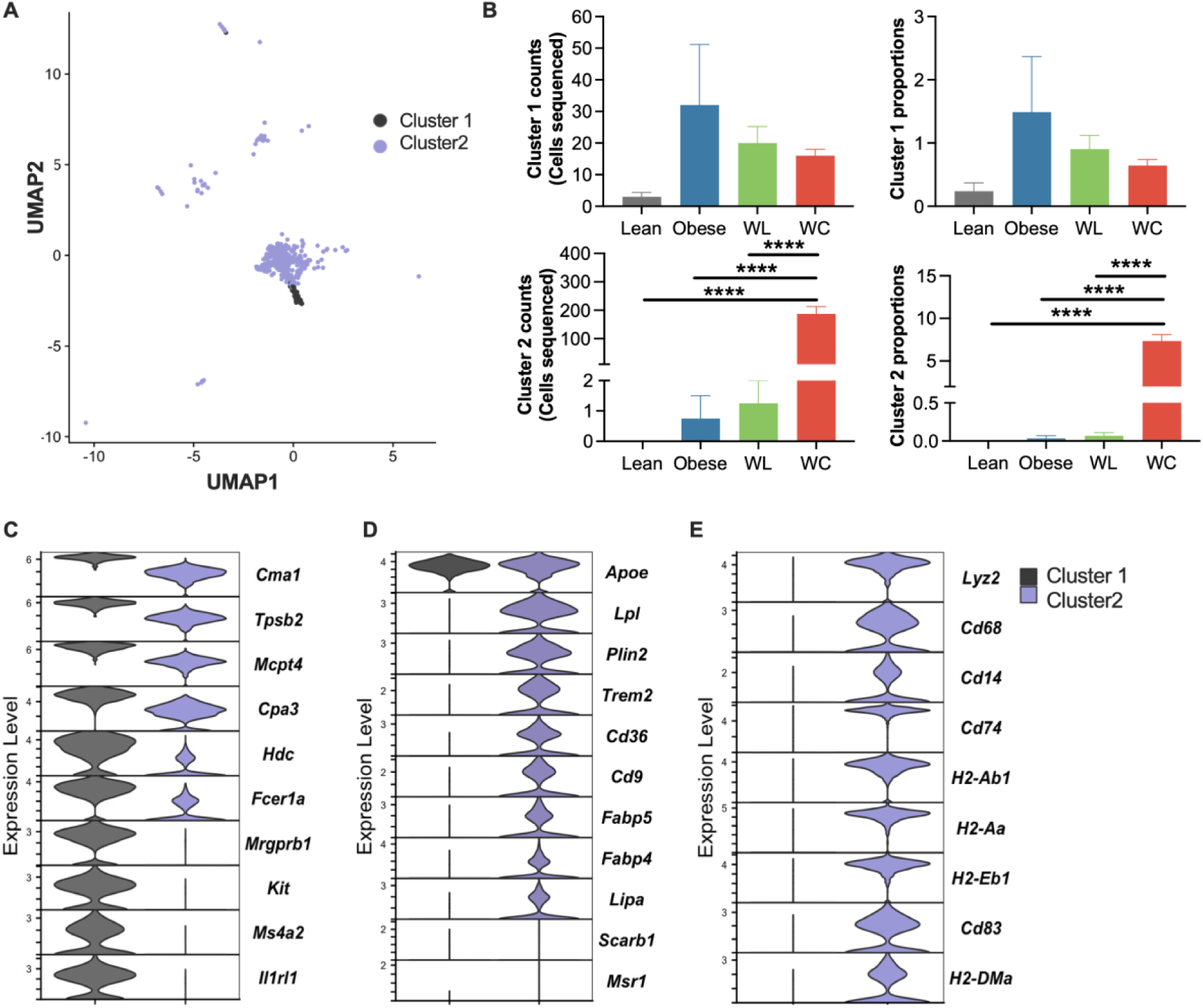
Weight cycling induces a distinct cluster of lipid handling mast cells. (A) UMAP of two distinct adipose mast cell clusters after subsetting on mast cells from [17]. (B) Mast cell counts (number of cells sequenced) and proportion (mast cells sequenced/ total cells) of both mast cell clusters. Violin plots of (C) mast cell and macrophage genes, as well as (D) lipid handling and antigen presentation related genes in each cluster. Expression level of all genes were significantly different between cluster 1 and cluster 2 by adjusted p-value with differential expression. All data were derived from new analyses of our previously published dataset [17].

### 3.4 Repeated exposure to adipose tissue conditioned media (ATCM) increases lipid handling in BMMC

Macrophages have a well-established role sensing and buffering lipids in the settings of obesity, atherosclerosis, and Alzheimer’s disease(17,38,39). However, it is currently not known if mast cells can acquire a similar lipid handling phenotype. Thus, to model the development of lipid-associated mast cells in culture, we cultured mature BMMC with ATCM. BMMC were treated ± ATCM for 24 hours, washed and cultured in growth media for 3 days, and then treated again ± ATCM for 24 hours (**Figure 4A**). ATCM had no impact on mast cell viability or differentiation (**Figure 4B**); however, repeated culture in ATCM did significantly increase mast cell lipid droplets as shown by Bodipy staining (**Figure 4C**). Repeated ATCM culture also significantly reduced the MFI of FcεR1 and cKIT on the surface of mast cells (**Figure 4D & E**). This data establishes a culture model by which repeated ATCM exposure increases lipid handling in mast cells while subsequently reducing expression of classical mast cell markers, as seen by scRNAseq from weight cycled mice.

**Figure 4:**
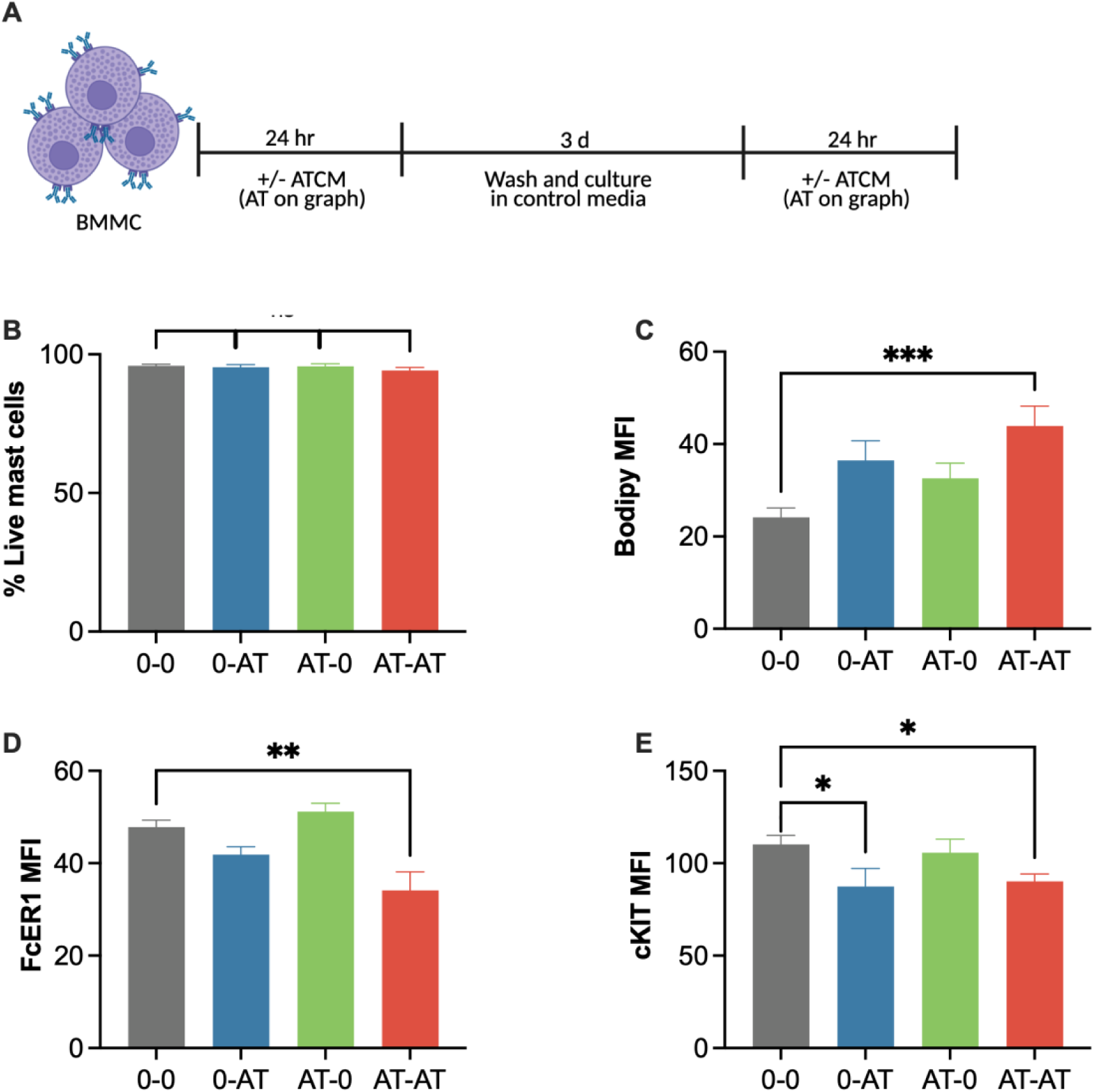
Repeated exposure to adipose tissue conditioned media (ATCM) induces lipid handling in culture. A) Schematic showing model of repeated exposure to ATCM (labeled AT on graph). B) Proportion of viable mast cells (cKit^+^ FcεR1^+^), C) Bodipy+ mast cells, D) mean fluorescence intensity (MFI) of FcεR1, and E) MFI of cKit at final timepoint by flow cytometry. Data are combined means ± SEM in n=2-4/ group from 3-6 independent experiments. *p<0.05, **p < 0.01, *** p <0.001.

### 3.5 Weight variance correlates with a cKit intermediate mast cell population in humans

Finally, to determine whether mast cell populations in humans are affected by weight cycling, we collected pilot data from four women with obesity and prediabetes. Immune cells were obtained from the subcutaneous adipose tissue and weight history was collected via questionnaire. Baseline participant characteristics include age 56.3±12.0 years, weight 109.6±27.7 kg, BMI 40.9±9.6 kg/m2, and fasting glucose 102.5±5.4 mg/dL. Using FMO controls, we gated on cKit^+^ FcεR1^+^ cells in the adipose tissue samples (**Figure 5A**). Interestingly, there appeared to be two diverse populations of mast cells, one with very high cKit expression (cKit^hi^ FcεR1^+^) and one with more intermediate cKit expression (cKit^int^ FcεR1^+^). The cKit^int^ FcεR1^+^ population was more prevalent than the cKit^hi^ FcεR1^+^ population (**Figure 5B**). The cKit^hi^ FcεR1^+^ population had increased surface expression of CD9 by MFI (**Figure 5C**). By comparison, the cKit^int^ FcεR1^+^ population had a greater percentage of CD68^+^ cells and greater CD68 surface expression by MFI. Additionally, there was a trend towards increased MFI of CD36 in the cKit^int^ FcεR1^+^ cells (**Figure 5D**).

**Figure 5:**
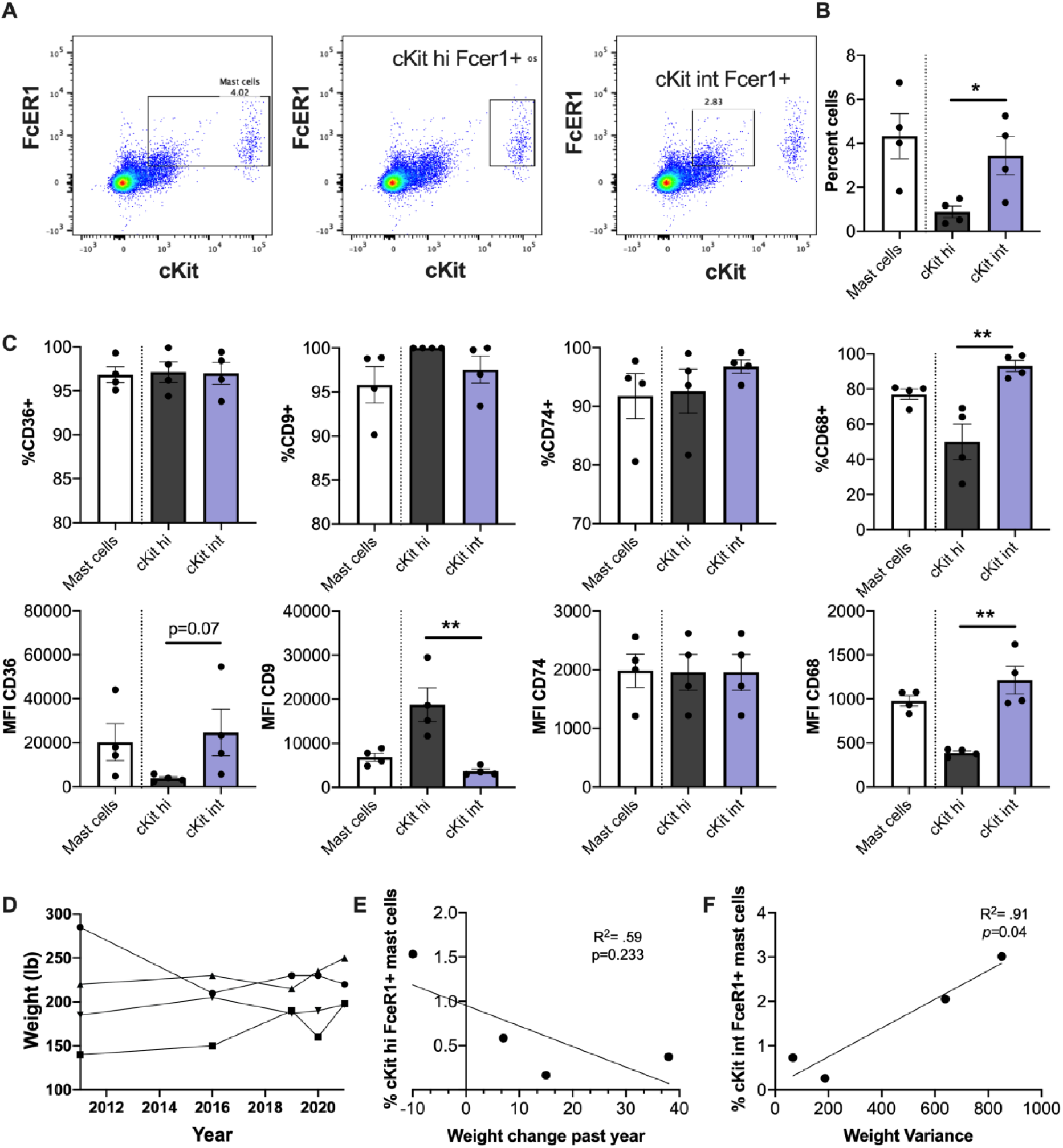
Weight variability correlates cKit intermediate mast cells in humans. (A) Representative flow plots for cKit^+^ FcεR1^+^, cKit^hi^ FcεR1^+^, and cKit^int^ FcεR1^+^ mast cell populations from human stromal vascular fraction. (B) Quantification of each mast cell population. (C) Quantification of percent of mast cells and MFI for the lipid handling receptors CD36 and CD9 and the antigen presentation proteins CD74 and CD68 on each mast cell population by flow cytometry. (D) Weight change plotted over time for each subject. (E) Correlation between the weight change in the previous year with the % cKit^hi^ FcεR1^+^ cells. (F) Correlation between weight variance and the % cKit^int^ FcεR1^+^ mast cells. Data are means ± SEM in n=4. *p<0.05, **p < 0.01.

We next determined whether these mast cell populations were different based on the weight history of the four study participants (**Figure 5D**). The participant who had most recently lost weight had the greatest population of the cKit^hi^ FcεR1^+^ cells, although there was no significant relationship between weight change in the past year and cKit^hi^ FcεR1^+^cells (**Figure 5E**). To examine weight cycling, we calculated weight variance (standard deviation squared) in all participants. There was a significant correlation between weight variance and the cKit^int^ FcεR1^+^ cells (**Figure 5F**). These data suggest that women who weight cycle have adipose mast cells with a different expression pattern of mast cell identity markers, lipid handling, and antigen presentation related surface proteins.

## 4. Discussion

Many studies show that immune cell populations change with adipose tissue expansion(40–42). However, despite the high prevalence of weight loss and weight regain in the US population, only a few studies measure the proportion and function of adipose immune cells in these conditions(19,41,43–46). Our data suggest that the prevalence of adipose mast cells change with weight loss and weight cycling, and we identify a novel population of lipid-associated mast cells that we can replicate in culture and identify in a small human pilot cohort.

Using new scRNAseq analysis, we identified two distinct mast cell clusters in the adipose tissue of male mice. The first is a cluster found in all mice that highly expresses the canonical mast cell surface receptors and proteases such as *Fcer1, Kit, Cma1,* and *Tpsb2* among many others. We consider these the classical mast cell population.

Additionally, there is a second cluster of cells that also express classical mast cell markers at a high level relative to all other immune cells, but at a lower level compared to the classical mast cell cluster. These cells also have increased expression of genes that play a role in lipid handling, lysosomal processing, scavenging, and antigen presentation. Interestingly, many of these genes are also elevated in a cluster of adipose macrophages following weight gain, first identified and named “lipid associated macrophages”, or LAMs, in 2019(17). In our dataset, the LAMs and lipid-associated mast cells cluster nearby on the UMAP, but are two distinct groups based on marker expression(19). Importantly, the macrophage populations do not express mast cell genes and the lipid-associated mast cells still have evident (though reduced) expression level of mast cell associated genes, suggesting a novel population.

To note, technical limitations restricted our ability to detect two populations of adipose mast cells using conventional techniques like flow cytometry or Toluidine blue staining in our murine groups. One limitation is that collagenase used for tissue dissociation can cleave the mast cell surface proteins (**Supplemental Figure 5**). Surface marker cleavage is also one reason we believe there is very limited data reported on murine adipose mast cells by flow cytometry (to our knowledge, this is only the second paper to attempt to quantify murine adipose mast cells by flow cytometry(47)). There is also substantial autofluorescence in adipose samples(48), contributing to measurement noise, especially in lipid dense cells. Thus, we developed a cell culture model to induce lipid handling in bone marrow derived mast cells as evidence that mast cells themselves can acquire a lipid handling phenotype. Interestingly, this was accompanied by reduced cKIT and FcεR1 protein expression, like our gene signature by sequencing. Interestingly, Jesse William’s group has shown that TREM2 agonist treatment can induce a lipid associated macrophage-like signature in stromal epithelial cells in a model of atherosclerosis(38), again supporting the induction of this cellular profile in non-macrophage cells. While flow cytometry did not identify a higher proportion of mast cells, CD63 (LAMP3) was increased in weight gain and weight cycled mice. We initially measured CD63 as a marker of degranulation, as it can be found on mast cell granules fusing at the surface during degranulation, however it also seems to be a lysosomal component that can sort cholesterol into intraluminal vesicles and contribute to lipid droplet accumulation(49–51). Thus, the change in expression may contribute to changes in lipid handling more than degranulation in our studies.

Only one other group has previously shown the presence of lipid droplets in mast cells following 6 days of insulin treatment(52,53). However, the specific expression of these lipid handling genes was not reported in the studies and little work has been done to show a mechanism beyond ER stress. Moreover, we do not observe increased plasma insulin in weight cycled mice, and actually detect reduced insulin secretion from the pancreatic beta cells in response to maximal stimulation(36), suggesting that weight cycling may have a different effect from chronic hyperinsulinemia in our model. Future work should examine how different signals such as insulin, glucose, fatty acids, and leptin within the adipose environment and ATCM affect lipid breakdown, storage, and biosynthesis in mast cells. In macrophages, activation with signals like LPS or IFNγ increase triglyceride storage and lipid droplet formation(54,55), and macrophages from obese fat upregulate both lipid droplet storage and lysosomal degradation of lipids(56). LAM development is also dependent upon TREM2(17,57,58), and thus, mechanisms of lipid scavenging, metabolism, and signaling should be further explored in our model. Last, further functional analyses should be used to identify the different lipid and inflammatory profiling of lipid-associated mast cells both *in vitro* and *ex vivo*.

Excitingly, we did find two adipose mast cell populations in subcutaneous adipose samples from obese women tested in our pilot work, which increases the potential translation of our findings. The population of cells with intermediate cKit expression was significantly associated with the weight variance of these subjects, like the lipid-associated mast cells observed by scRNAseq. Moreover, this population had higher expression of CD68, and a trend towards increased CD36, which was also observed in the scRNAseq data. However, there was lower expression of CD9 in the cKit intermediate mast cells than the population with higher Kit expression. We should emphasize that our murine data was primarily collected from male epididymal adipose tissue while our human data was collected from female subcutaneous adipose tissue. Moreover, our body weight history was self-reported. Thus, future studies should further assess differences by sex, adipose depots, and/or organism and should use additional data collection measures like electronic medical records when feasible. However, in support of our data, a study by Lopez-Perez et al. also showed a change in adipose mast cells in adults with obesity and type 2 diabetes. In their first cohort, a reduction in classical adipose mast cells from omental fat was observed(59). Upon follow-up, the authors noticed that omental adipose mast cells had reduced surface expression of CD45, cKit, and FcεR1(60). However, it is not clear if they also observed an intermediate population in their participants or if the effects of weight cycling are required to induce this intermediate population.

Together, our data identify a novel lipid-associated mast cell population. Future studies should not only further explore the systemic impact of mast cells in obesity and weight cycling, but the specific function of mast cells based on polarization and/or lipid handling. Moreover, it would be exciting to explore what other homeostatic or disease contexts could give rise to lipid-associated mast cell populations. In sum, lipid-associated mast cells may pose a novel contribution to adipose metabolism in weight cycling in both mice and humans.

## Authorship

MK and AMB assisted with cell culture model development, data collection, analysis and manuscript writing/editing. MAC assisted with sequencing data collection, analysis, and interpretation. HJS provided the weight cycling questionnaire and MM provided the human samples. Both assisted with study design and data analysis. AHH assisted with study design and data interpretation and provided funding for the murine cohort data. HLC conceptualized the study, obtained and analyzed the data, drafted the manuscript, provided funding for the cell culture work and human sample analysis, and is the guarantor of the work. All authors contributed to manuscript revisions.

## Acknowledgements

This project was funded by a Veterans Affairs Merit Award 5I01BX002195 to AHH. AHH was also supported by a Career Scientist Award from the Veterans Affairs (IK6 BX005649). HLC was funded by an American Heart Association Postdoctoral Fellowship (20POST35120547) and is currently funded by an American Heart Association Transformational Project Award (25TPA1471583). MAC was funded by an NIH F31 Predoctoral Fellowship (1F31DK123881), and both MAC and HLC were previously supported by the Molecular Endocrinology Training Program (T32 DK07563). Additionally, the human mast cell work was funded by The Vanderbilt Institute for Clinical and Translational Research (VICTR) grants to HLC and MM, a NIH KL2 (KL2TR002245) and K23 (K23HL159351) to MM, and utilized resources of the Tennessee Center for AIDS Research (P30AI110527). VICTR is funded by the National Center for Advancing Translational Sciences (NCATS) Clinical Translational Science Award (CTSA) Program, Award Number 5UL1TR002243-03. Figure schematics were created with Biorender.com.

---

**Supplemental Figure 1:**
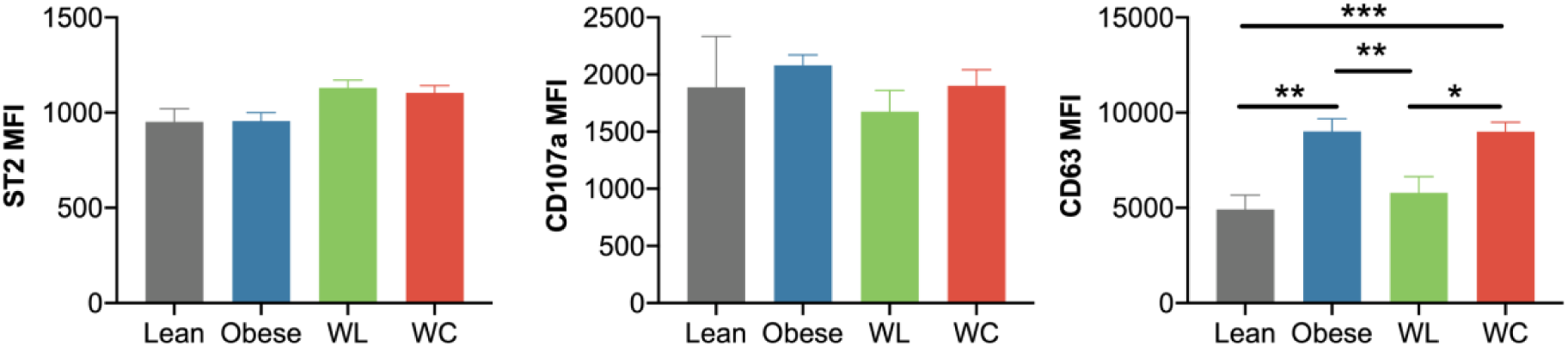
CD63 is increased in mast cells from obese and weight cycled mice. MFI of ST2, CD107a, and CD63 on epididymal adipose mast cells by flow cytometry. Data are means ± SEM in n= 8-12/ group. *p<0.05

**Supplemental Figure 2:**
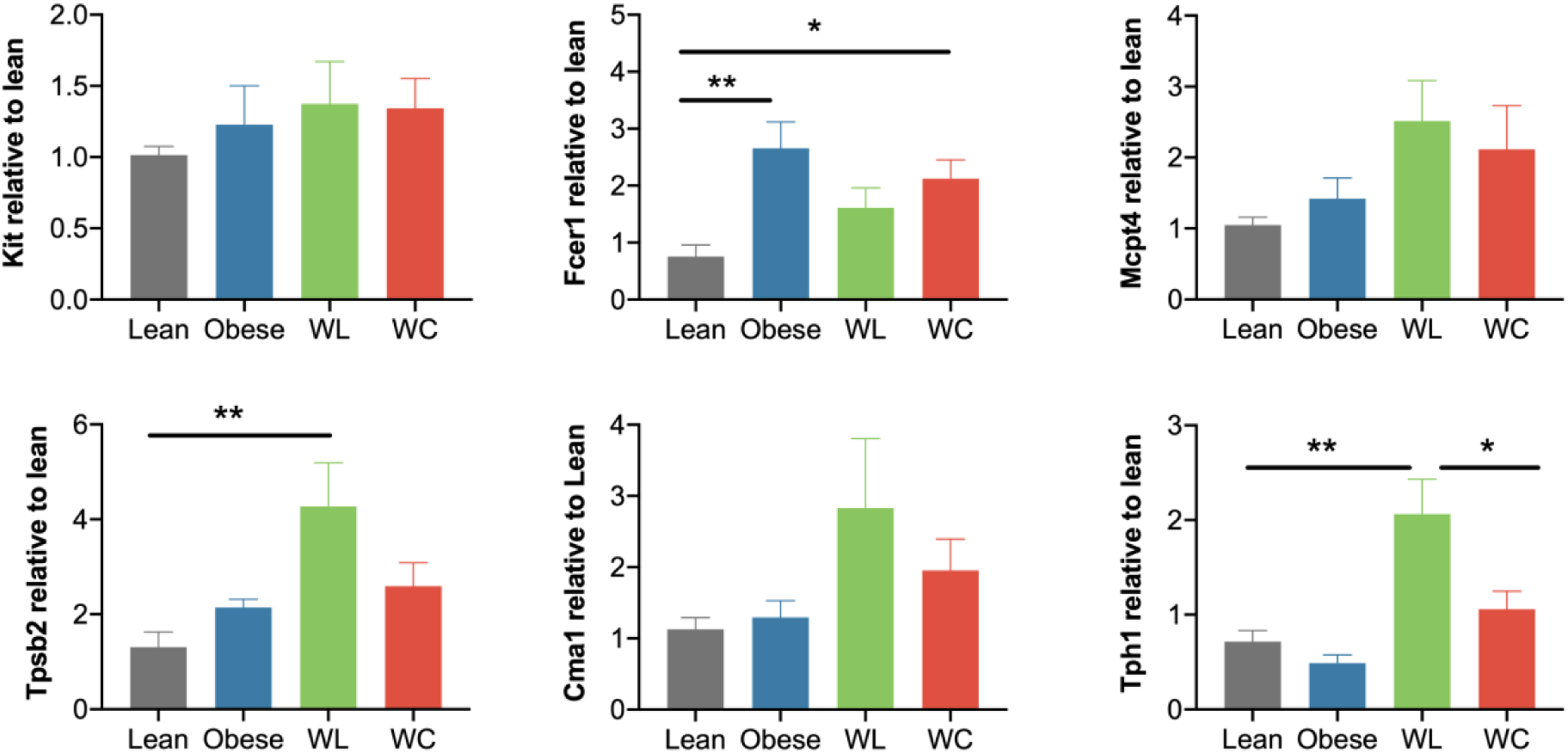
Weight cycling does not increase mast cell gene expression beyond that of the obese group. Expression of mast cell associated genes from the subcutaneous adipose tissue by qPCR (n=6-10/group).

**Supplemental Figure 3:**
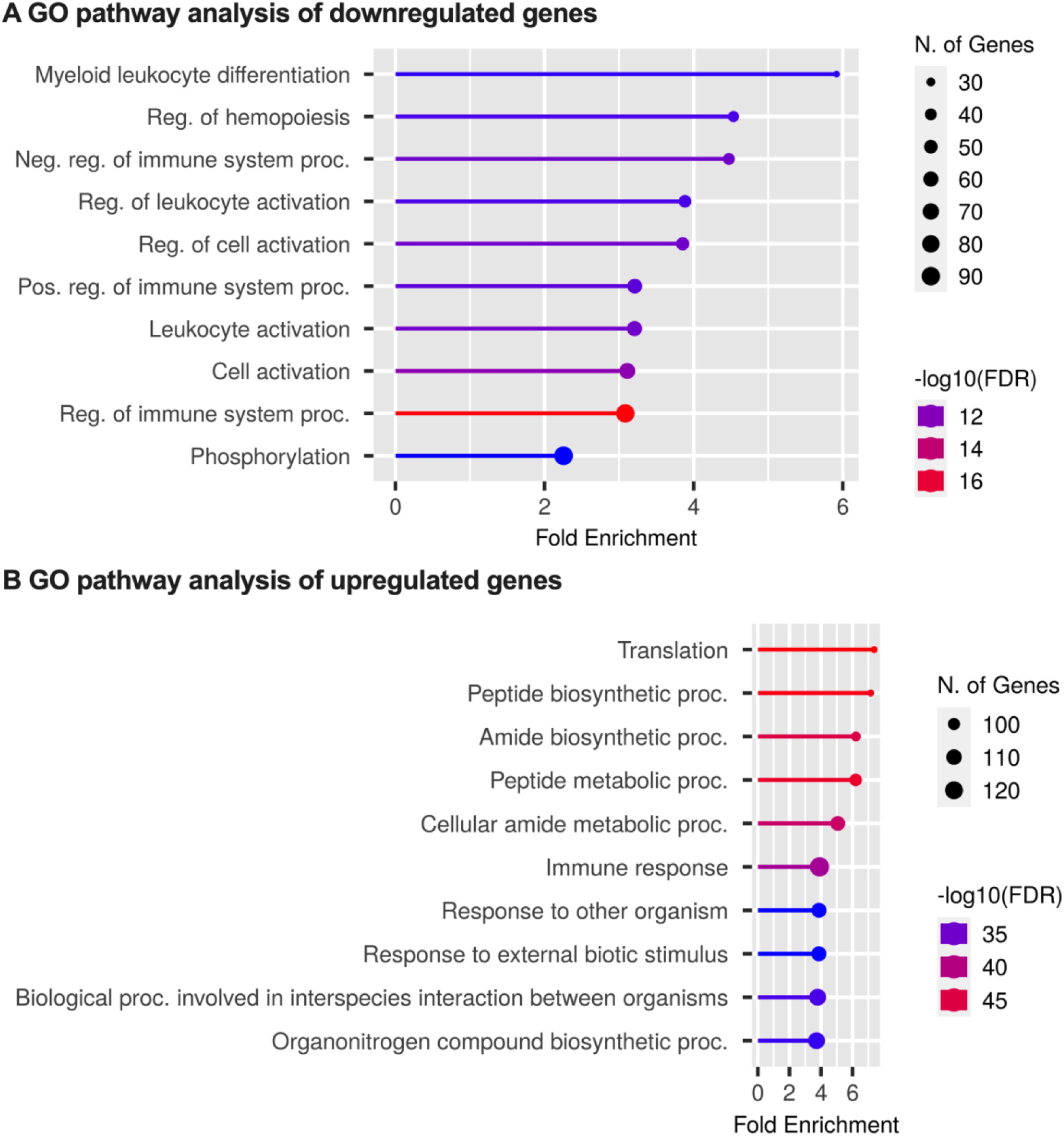
Pathway analysis of differentially expressed genes in new mast cell cluster. GO pathway analysis of significantly (A) downregulated and (B) upregulated genes in cluster 2 vs. cluster 1 generated from our previously published dataset [17].

**Supplemental Figure 4:**
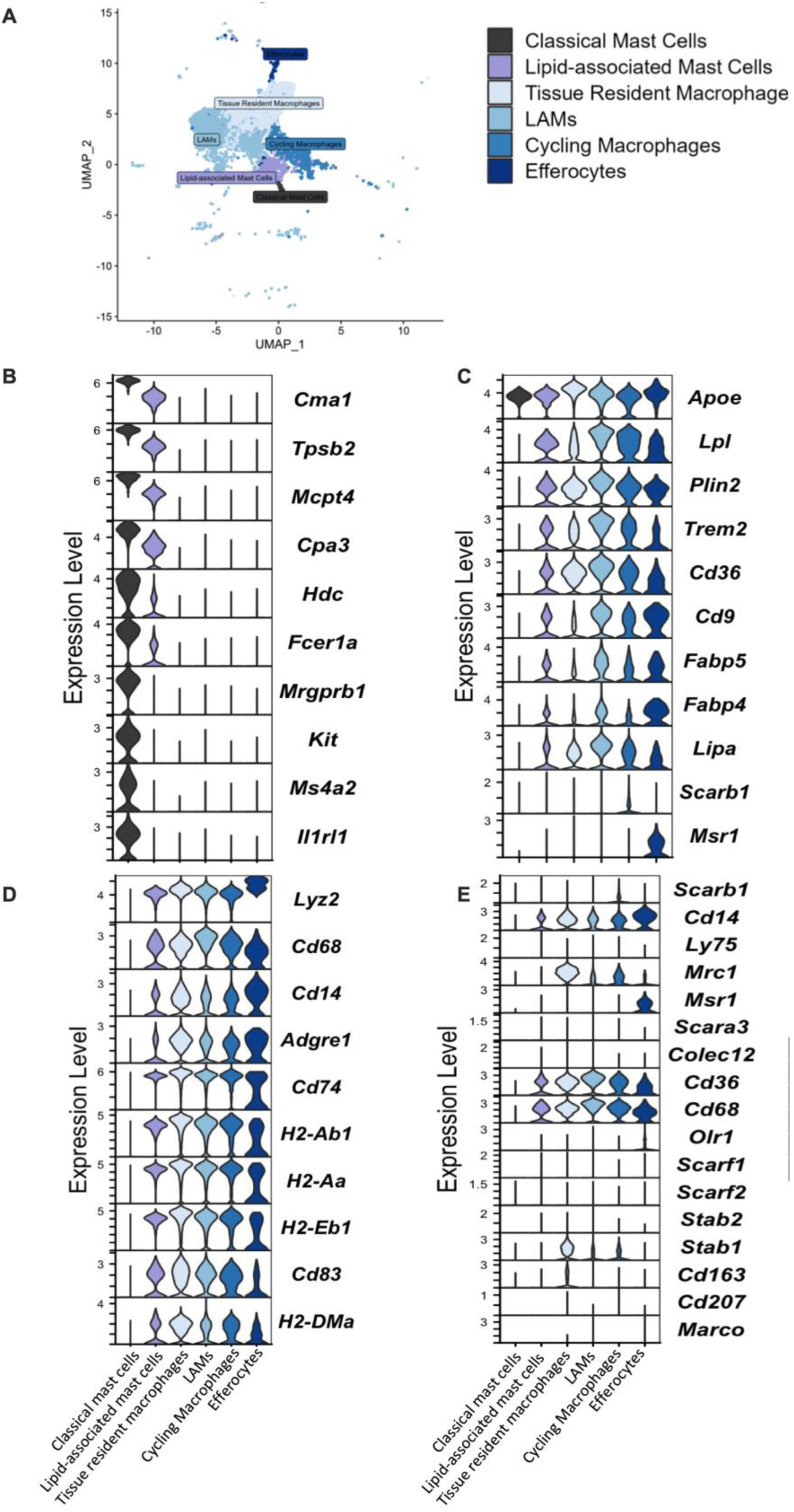
UMAP clustering and violin plots for gene expression between macrophage and mast cell subclusters. (A) UMAP of the mast cell and macrophage subclusters. Violin plots of (B) mast cell, (C) lipid handling, (D) lysosomal and antigen presentation, and (E) scavenger receptor genes among all mast cell and macrophage subclusters generated from our previously published dataset [17].

**Supplemental Figure 5:**
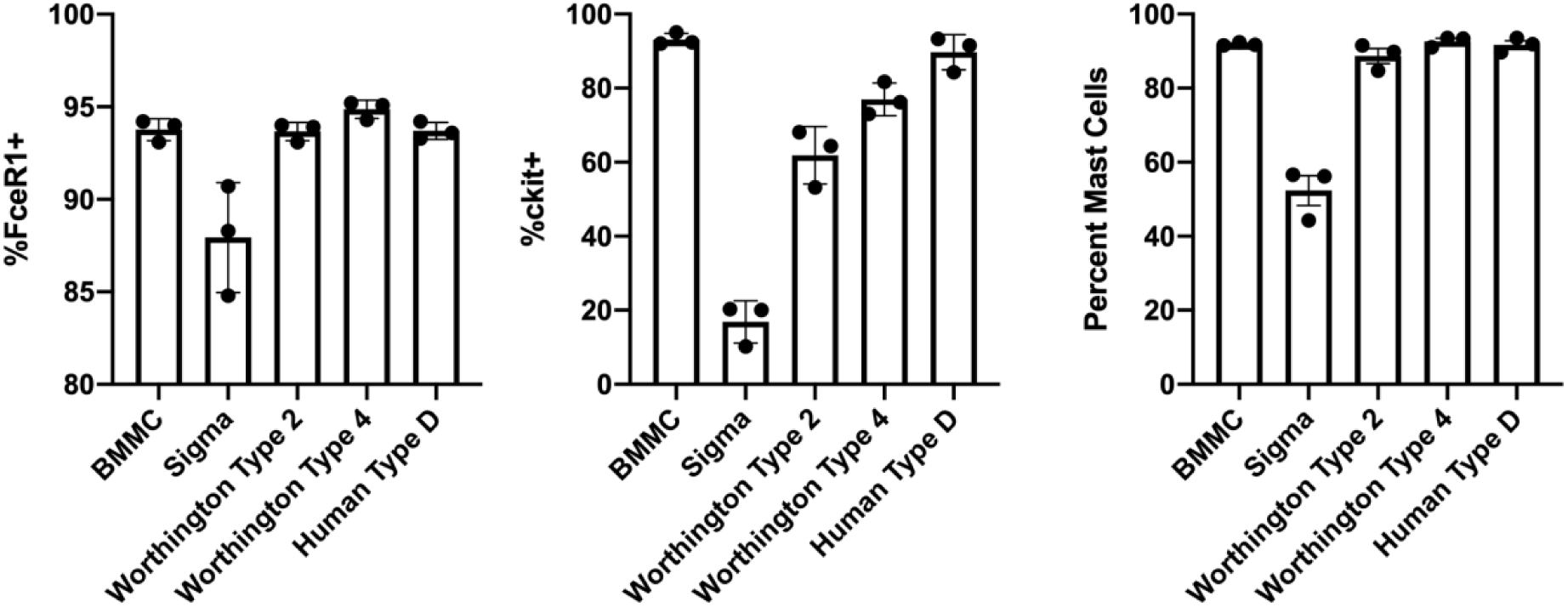
Collagenase cleaves mast cell surface markers. Flow cytometry analysis for cKit and FcER1 of BMMC treated with 2 mg/ mL Sigma type IV, Worthington type II or IV, or Roche type D collagenase for 30 min at 37°C.

